# Prospects for a sequence-based taxonomy of influenza A virus subtypes

**DOI:** 10.1101/2023.07.06.548035

**Authors:** Art F. Y. Poon

## Abstract

The hemagglutinin (HA) and neuraminidase (NA) proteins are the primary antigenic targets of influenza A virus (IAV) infections. IAV infections are generally classified into subtypes of HA and NA proteins, *e.g.*, H3N2. Most of the known subtypes were originally defined by a lack of antibody cross-reactivity. However, genetic sequencing has played an increasingly important role in characterizing the evolving diversity of IAV. Novel subtypes have recently been described solely by their genetic sequences, and IAV infections are routinely subtyped by molecular assays, *e.g.*, real-time PCR, or the comparison of sequences to references. In this study, I carry out a phylogenetic analysis of all available IAV protein sequences in the Genbank database (over 1.1 million records) to determine whether the serologically-defined subtypes can be reproduced with sequence-based criteria. I show that a robust genetic taxonomy of HA and NA subtypes can be obtained with a simple clustering method, namely by progressively partitioning the phylogeny on its longest internal branches. However, this taxonomy also requires some amendments to the current nomenclature. For example, two IAV isolates from bats previously characterized as a divergent lineage of H9N2 should be separated into their own subtype. With the exception of these small and highly divergent lineages, the phylogenies relating each of the other six genomic segments do not support partitions into major subtypes.

## Introduction

Influenza A virus (IAV) genomes are comprised of eight single-stranded, negative sense RNA segments, which are numbered in decreasing order with respect to their lengths. The diversity of IAV is usually characterized by the two segments (numbered 4 and 6) carrying the genes encoding the hemagglutinin (HA) and neuraminidase (NA) proteins, respectively. HA and NA are surface-exposed proteins responsible for binding and release of virus particles from host cells, respectively. As significant antigenic targets of the adaptive immune system, they are the most rapidly evolving proteins of the IAV genome [3]. The global diversities of both HA and NA sequences are clustered into serotypes or subtypes, terms which are used interchangeably in the literature; herein I will use the term ‘subtypes’ for consistency. These subtypes are identified by H and N prefixes for the respective proteins, followed by an integer suffix, producing the widely familiar HnNn nomenclature, *e.g.*, H3N2. Certain combinations of HA and NA subtypes are observed more frequently than others due to pandemic growth. Although these combinations are also referred to as subtypes, each genome segment has its own subtype nomenclature.

There are presently 18 described HA subtypes that are numbered H1 to H18 (Table 1). In addition, a potential new subtype H19 has only recently been described [7] from a partial HA sequence isolated in 2008 from waterfowl in Kazakhstan. In addition, there are 11 described subtypes of NA (N1 to N11). These subtypes were originally defined solely from their antigenic characteristics, predominantly from hemagglutination inhibition (HI, specifically for HA), complement fixation (CF), or double immunodefusion (DID [5]) testing. For instance, H2, initially labelled the ‘Far East’ or ‘Asian’ strain of IAV [6], was distinguished from H1 by the lack of HI activity by sera from hosts infected by the other subtype [14]. The nomenclature for HA and NA subtypes also formerly incorporated the host organism [34]. For instance, H1, Hsw1, Heq1 and Hav1 were originally different HA subtypes isolated from human, swine, equine and avian hosts, respectively [34]. These were subsequently merged by Schild et al. [26] into the current host-agnostic Hn and Nn nomenclature on the basis of the cross-reactivity of isolates from different host species.

**Table 1:** Summary of the currently known HA subtypes and the methods used to describe them. Previous subtype and prototype strain designations for subtypes H1 to H12 were obtained from Schild et al. [26]. Methods: HI = hemagglutination inhibition; CF = complement-fixation; DID = double immunodiffusion; P = phylogeny; AAI = amino acid sequence identity. *The authors use an immunodiffusion assay that resembles DID (described in Beard [2]). †H19 has only recently been proposed from a single partial HA sequence.

| Subtype | Previous designation | Prototype strain | Methods | Publication |
| --- | --- | --- | --- | --- |
| H1 | H0 | A/Puerto Rico/8/1934 | HI, DID | [26] |
|  | H1 | A/Fort Monmouth/1/1947 |  |  |
|  | Hsv1 | A/Swine/Wisconsin/15/30 |  |  |
| H2 | H2 | A/Singapore/1/57 | HI | [14] |
| H3 | H3 | A/Hong Kong/1/68 | HI, DID | [26] |
|  | Heq2 | A/eq/Miami/1/63 |  |  |
|  | Hav7 | A/Duck/Ukraine/1/63 |  |  |
| H4 | Hav4 | A/Duck/Czech/56 | HI, CF, DID | [21] |
| H5 | Hav5 | A/Tern/South Africa/61 | HI, CF | [21] |
| H6 | Hav6 | A/Turkey/Mass/3740/65 | HI, CF | [21] |
| H7 | Heq1 | A/Equine/Prague/1/56 | HI, DID | [26] |
|  | Hav1 | A/FPV/Dutch/27 |  |  |
| H8 | Hav8 | A/Turkey/Ontario/6118/68 | HI, CF, DID* | [18] |
| H9 | Hav9 | A/Turkey/Wisconsin/66 | HI, DID | [31] |
| H10 | Hav2 | A/Chick/Germ/N/49 | HI, CF | [21] |
| H11 | Hav3 | A/Duck/England/56 | HI, CF | [21] |
| H12 | Hav10 | A/Duck/Alberta/60/76 | HI, DID | [12] |
| H13 |  | A/Gull/Md/704/77 | HI, DID, AAI | [13] |
| H14 |  | A/Mallard/Gurjev/263/82 | HI, DID, P, AAI | [17] |
| H15 |  | A/duck/Australia/341/83 | HI, DID, P, AAI | [24] |
| H16 |  | A/black-headed gull/Sweden/5/99 | HI, DID, P, AAI | [8] |
| H17 |  | A/little yellow-shouldered bat/Guatemala/153/2009 | P, AAI | [29] |
| H18 |  | A/flat-faced bat/Peru/033/2010 | P, AAI | [30] |
| H19† |  | A/Common Pochard/Kazakhstan/KZ-52/2008 | P, AAI | [7] |

Since the description of subtype H13 by Hinshaw et al. [13] in 1983, genetic sequence analysis has been increasingly incorporated into the characterization of novel HA and NA subtypes (Table 1). Indeed, the last three HA subtypes (H17-H19) have been described solely by their genetic sequences [7, 29, 30]. High-throughput genetic sequencing has become a ubiquitous feature of clinical and public health laboratories, boosted recently by the expansion of resources to confront the global SARS-CoV-2 pandemic. For instance, the number of IAV sequences deposited in the GISAID (Global Initiative for Sharing All Influenza Data, https://gisaid.org) database has grown exponentially from 943 sequences in 1992 to 307,198 in 2022 (Supplementary Figure S1).

Here, I investigate whether it is feasible to distinguish HA and NA subtypes based only on genetic diversity. This is a different problem than the supervised classification of genetic sequences to subtypes based on their similarity to a predefined set of reference sequences, commonly performed by BLAST search [27]. Instead, I ask whether an unsupervised method can robustly cluster HA and NA sequences into the antigenically-defined subtypes. Furthermore, I assess whether it is feasible to apply a similar classification scheme to the other six genomic segments. There is surpisingly limited work in this area. When describing serotype H14, for instance, Kawaoka et al. [17] proposed that HA sequences that differed at 30% of amino acids or more should be separated into different subtypes. However, this proportional (p) distance criterion is not attained for all pairwise comparisons of subtypes [24, 29]. This method also makes no adjustment for multiple substitutions at the same sites over long evolutionary time scales. Furthermore, the shortest p-distance between groups can depend by chance on which sequences have been incorporated into the analysis. Ideally the phylogenetic analysis should incorporate as many genetic sequences as possible.

## Methods

I downloaded all available protein sequences associated with all eight IAV segments from the NCBI Genbank database (accessed June 6, 2023 except for HA, which were retrieved April 28, 2023), using a minimum length filter to exclude records with incomplete nucleotide sequences (Supplementary Table S1). For each segment, I used regular expressions to filter sequences from other segments based on gene and protein annotations in the sequence names. In addition, I excluded sequences derived from alternate open reading frames (PB1-F2 and PA-X), and concate-nated sequences involved in alternative splicing with overlapping regions excluded (M1/M2 and NS1/NS2). Next, I discarded sequences with more than 10% ambiguous residues (‘X’) and compressed the remainder into unqiue sequences, recording the labels of duplicate sequences into a separate file. The remaining sequences were aligned using MAFFT (version 7.453) [16]. I manually evaluated and amended the resulting alignments with AliView (v.2018) [19]. Using FastTree, I reconstructed a preliminary tree from the alignment without maximum-likelihood or minimum-evolution steps (*i.e.*, neighbor-joining), and then visually assessed the tree using Taxonium [25]. After excluding any problematic sequences identified in the preliminary tree (*e.g.*, recombinant sequences produced by molecular cloning [11]), I re-ran FastTree with the default optimization steps to generate a maximum likelihood tree.

I investigated two different methods to partition each tree into monophyletic clades (*i.e.*, subtrees) as putative subtypes. First, I calculated the following quantities for each internal node in the tree:

1. *y_i_*, the mean tip-to-tip (patristic) distance between every tip descending from node *i*;
2. *d_i_*, the mean distance from node *i* to every descendant tip; and
3. *v_i_*, the total distance from node *i* to its sibling node *j*.

The quantities *d_i_* and *y_i_* were calculated by postorder traversal of the tree for increased efficiency. For instance, *y_i_* was calculated by this recurrence relation, where *a* and *b* are child nodes of *i*:

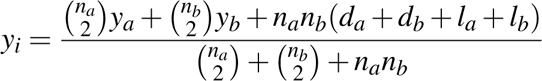

where *n_a_* is the number of tips descending from node *a*, and *l_a_* is the length of the branch associated with (below) node *a*. Tip nodes were initialized with *n_i_* = 1, *d_i_* = 0, and *y_i_* = 0. Hence, the *y_i_*for the internal node of a cherry reduces to *l_a_* + *l_b_*. The quantity *y_i_* is analogous to the mean pairwise distance within a subtype, while *v_i_*is analogous to the shortest distance between two subtypes. Given minimum *v* and maximum *y* cutoffs, subtrees meeting these criteria were located by evaluating nodes by preorder traversal of the tree. This method will be referred to as ‘nodewise’ clustering.

Second, I progressively cut the tree into subtrees on internal branches (edges) with a length exceeding a given cutoff, using the following algorithm:

1. initialize subtree list *S* with input tree
2. for each subtree *s* in *S*

- locate longest internal branch *b_s_* in *s*
- if length of *b_s_* exceeds cutoff

- remove *b_s_* to yield subtrees *s*_1_ and *s*_2_
- for each new subtree, remove the node associated with *b_s_* and join the branches
- append *s*_1_ and *s*_2_ to new list *S^I^*
- otherwise append *s* to new list *S^I^*
3. replace *S* with *S^I^*
4. repeat from step 2 until no internal branches exceed cutoff

This method will be referred to as ‘edgewise’ clustering. Both algorithms were implemented with the Phylo toolkit in Biopython [28].

To measure the correlation between the subtree clusters and the subtype annotations of sequences, I calculated the normalized mutual information (nMI) of these two partitions:

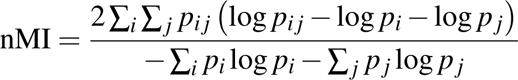

where *p_i_* is the marginal proportion of labeled tips in subtree *i*, *p _j_*is the marginal proportion of tips labeled with subtype *j*, and *p_i j_* is the number of tips in subtree *i* with label *j* divided by the total number of labels, *i.e.*, the joint proportion (see also Equation 14.4.17 in Press et al. [22]). The nMI adjusts for different numbers of subsets between these two partitions, and it ranges from 0 (no correlation) to 1 (for identical partitions).

## Results

Maximum likelihood trees for *n* = 66, 864 HA and *n* = 52, 146 NA amino acid sequences are displayed in Figure 1 with branches coloured by subtype annotations, *i.e.*, NCBI Genbank source qualifiers. As anticipated, the subtrees labelled by subtype annotations are distinctly separated by long internal branches in both trees, implying that these subtypes should be readily distingushed by basic sequence-based criteria. The overall scale of the NA tree is substantially greater than the HA, driven by the enormous divergence of N10 and N11 from the other subtypes (4.23 expected substitutions per site). Based on their discordant locations in the tree, I identified 15 HA and 30 NA sequences with incorrect subtype annotations (Supplementary Tables S2 and S3). In addition, I classified 2,353 HA and 2,563 NA sequences with missing subtype annotations (*e.g.*, ‘mixed’, ‘HX’ or ‘unknown’) by locating their nearest neighbours in the respective phylogenies. These sequences were substantially less likely to be assigned to the most common subtypes (Supplementary Figure S2). These inferred subtype labels were excluded from further analyses to mitigate potential biases.

**Figure 1:**
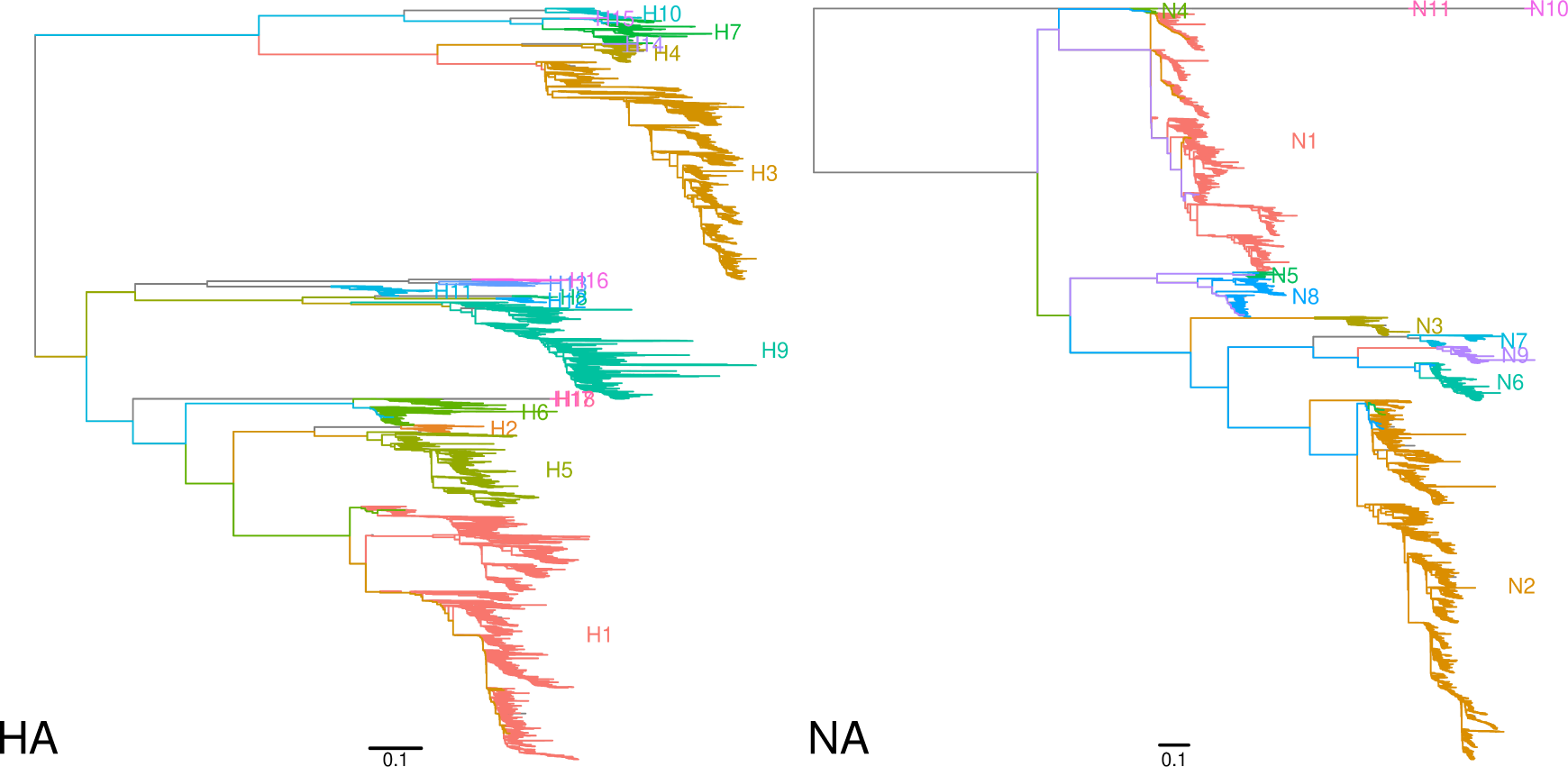
Maximum likelihood trees reconstructed from unique amino acid sequences of HA (*n* = 66, 864) and NA (*n* = 52, 146). The trees are midpoint-rooted and ladderized. Branches are coloured by respective ‘serotype’ annotations from the source qualifiers in the NCBI Genbank database. Scale bars indicate the branch length corresponding to 0.1 expected amino acid substituitons per site. These plots were generated using the R package *ggfree* (https://github.com/ArtPoon/ggfree).

I evaluated two different methods to partition a phylogeny into subtrees that are consistent with the antigenically-defined IAV hemagglutinin (HA) and neuraminidase (NA) subtypes. First, I assessed a nodewise method in which I calculated the mean tip-to-tip distance *y* and total branch length to the sibling node *v* for subtrees descending from every internal node. These quantites are analogous to the within- and between-group measures of genetic divergence that have been used to define clusters within IAV subtypes, *i.e.*, clades [1, 32]. Both *v* and *y* were required to partition the tree *de novo* into subtrees. For instance, using only a minimum cutoff on *v* tends to prematurely terminate a search by preorder traversal of nodes, *i.e.*, yielding two large subtrees that descend directly from the root node.

I evaluated the discordance between subtypes extracted from the sequence metadata (herein referred to as ‘labels’) and the phylogenetic subtrees produced at varying cutoffs of *y* and *v* (Supplementary Figure S3A-C). Overall, the discordance in the number of subtrees was minimized at a tip-to-tip distance cutoff of about *y* = 1.2. The results were less sensitive to varying cutoffs for *v*. However, the correspondence between subtrees and labels at the best settings was generally poor. Sequences labeled as subtype H9 were consistently partitioned into multiple subtrees. In addition, several groups of subtype labels become merged into single subtrees, namely H2/H5, H3/H4/H14, H7/H10, H8/H12, and H11/H13/H16 (Supplementary Figure S3D). These discrepancies could not be resolved by varying the clustering settings for this method.

Since the results of nodewise clustering on the HA phylogeny were so poor, I abandoned this method and implemented a simpler edgewise method. This method partitions the phylogeny by progressively cutting internal branches with lengths exceeding some cutoff. The number of subtrees obtained from the HA phylogeny under varying cutoffs and their correlation with subtype labels, as measured by normalized mutual information (nMI), are summarized in Figure 2A. Overall, the correlation between subtrees and subtype labels was high across a range of cutoffs, with nMI exceeding 0.99 at cutoffs from 0.12 to 0.2675, where nMI = 1 indicates perfect correlation. On the other hand, the expected number of 18 subtrees is attained only within a narrow range of cutoffs around 0.18. At this cutoff the partition of subtrees was highly concordant with subtype labels, as summarized in Figure 2B, with the exception of two discrepancies. First, one subtree (‘s8’) was comprised of only two sequences corresponding to IAV isolated from bats resembling H9 [15, 23], which predominantly circulates in avian host species. This subtree does not merge with the adjacent subtree (‘s7’) carrying the other 8,197 H9 labels until the cutoff is relaxed to 0.27, at which there is a sudden drop in the number of subtrees from 17 to 14 (Figure 2A). Second, the nine HA sequences labeled as H15 are grouped with H7 at this cutoff. They are split when the cutoff is decreased from 0.18 to 0.1775. If the cutoff is increased from 0.18 to 0.1825, then subtrees labeled H13 and H16 become merged into a single subtree.

**Figure 2:**
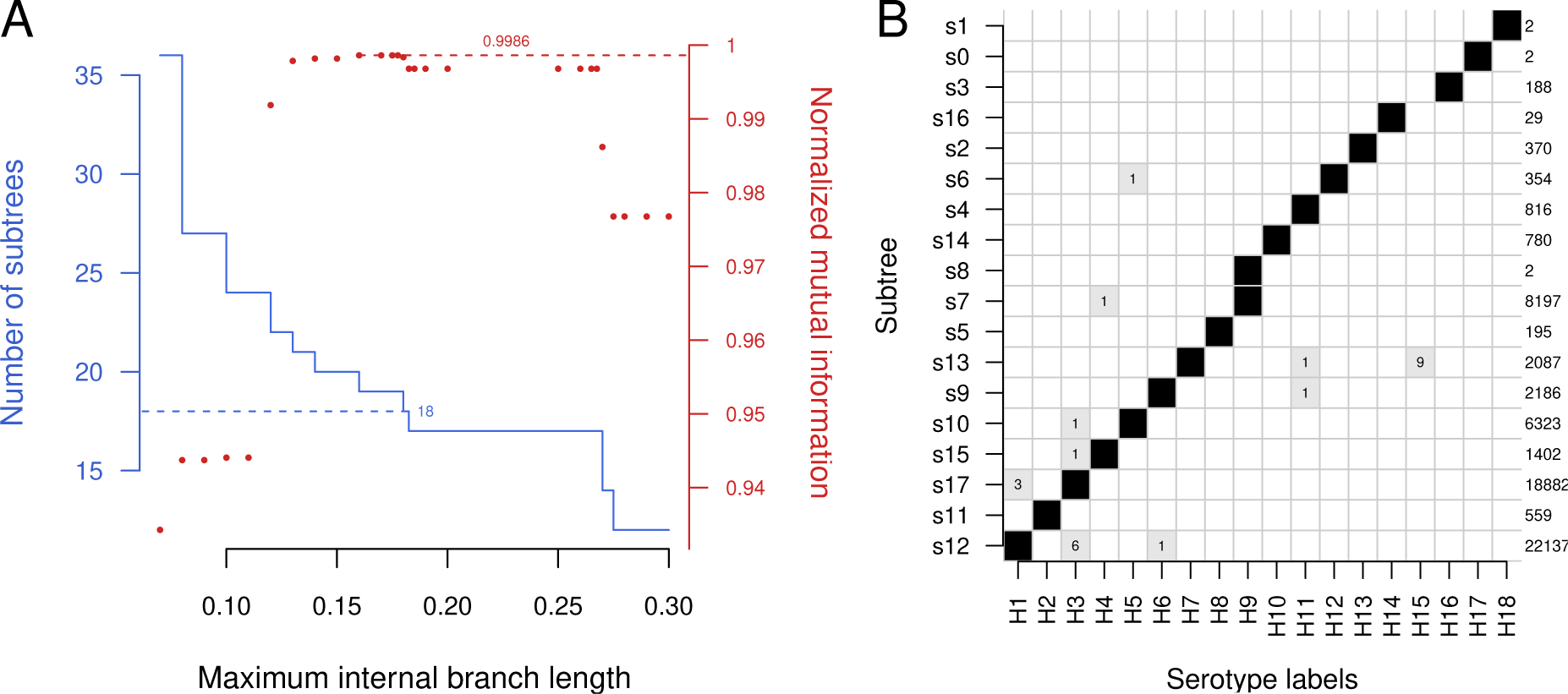
(A) A solid line (blue) indicates the number of subtrees produced by partitioning the phylogeny of HA sequences at varying branch length cutoffs (edgewise clustering). A dashed horizontal line is drawn at *n* = 18 to indicate the target number of subtrees. Points (red) indicate the normalized mutual information (nMI, measuring the correlation between subtree and HA subtype labels) at varying cutoffs. The maximum nMI at 99.86% is indicated by a dashed line. (B) Distribution of HA subtype labels among subtrees produced at an internal branch length cutoff of 0.18. Boxes are shaded in proportion to relative counts of labels. Total counts are listed along the right side. Individual counts are printed in the respective boxes for low-frequency labels, most of which correspond to misclassified sequences (see Table S2).

Next, I applied the edgewise method to a maximum-likelihood phylogeny of 52, 146 NA se-quences. This method yielded 12 subtrees for a broad range of cutoffs (from 0.222 to 0.4 expected substitutions, Figure 3A). Similarly to the case with the HA phylogeny, the target number of 11 subtrees could only be obtained within a narrow range of cutoffs (from 0.405 to 0.425). At a cutoff of 0.41, I observed the same two major discrepancies among the resulting 11 subtrees (Figure 3B). Two sequences labeled as N2 were separated from the subtree carrying the bulk of the remaining N2 labels (*n* = 23, 803). Not surprisingly, these corresponded to the same divergent H9N2-like viruses isolated from bats as noted above [15, 23]. In addition, 394 sequences labeled as N5 are placed in the same subtree as N8 unless the cutoff is reduced below 0.405.

**Figure 3:**
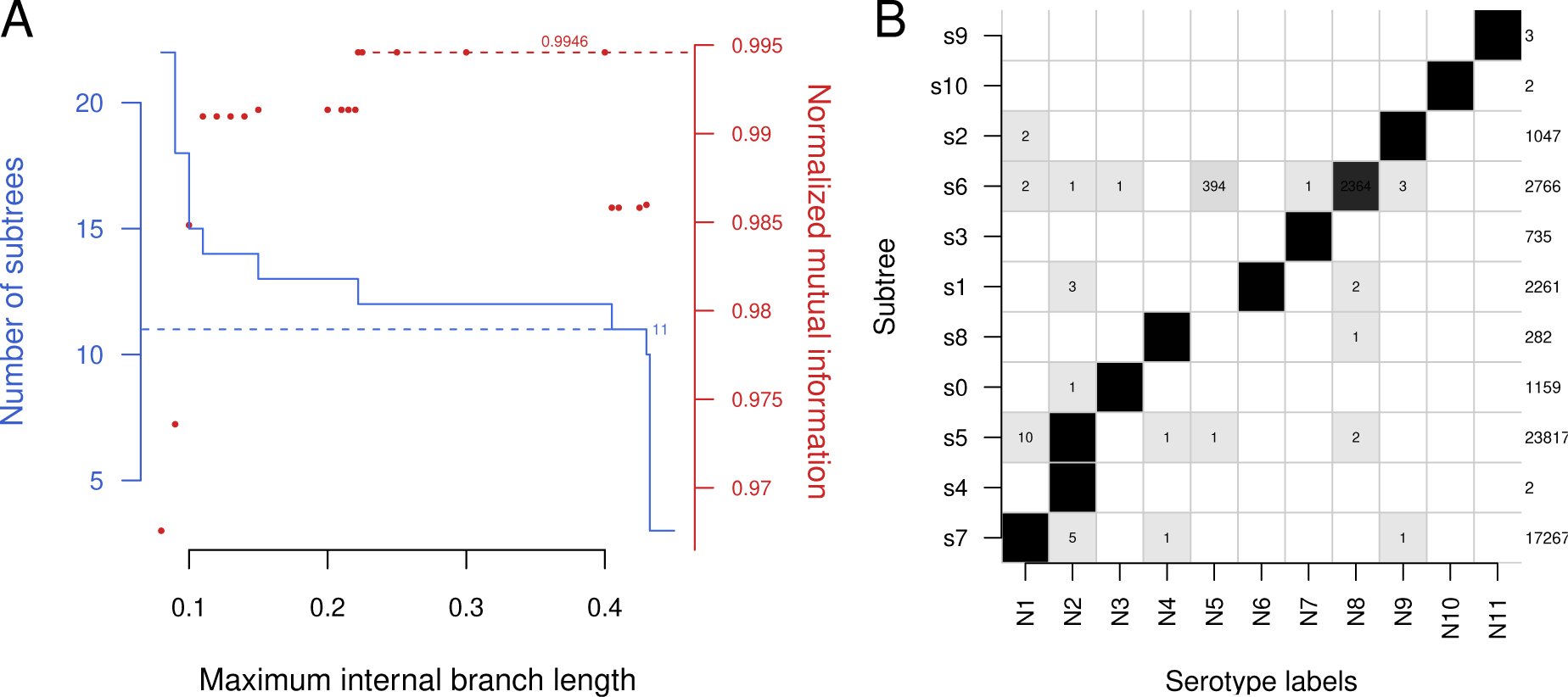
(A) A solid line (blue) indicates the number of subtrees produced by partitioning the phylogeny of NA sequences at varying branch length cutoffs (edgewise clustering). A dashed horizontal line is drawn at *n* = 11 to indicate the target number of subtrees. Points (red) indicate the normalized mutual information (nMI, measuring the correlation between subtree and NA subtype labels) at varying cutoffs. The maximum nMI at 99.46% is indicated by a dashed line. (B) Distribution of NA subtype labels among subtrees produced at an internal branch length cutoff of 0.41. Boxes are shaded in proportion to relative counts of labels. Total counts are listed along the right side. Individual counts are printed in the respective boxes for minority serotypes. The smaller counts generally correspond to misclassified sequences (see Table S3).

Finally, I repeated this same process for the remaining IAV genome segments. To increase the amount of genetic variation available for reconstructing trees, I concatenated the non-overlapping regions of the M1 and M2 proteins encoded on segment 8, and the NS1 and NS2 proteins encoded on segment 8. Unlike the HA and NA phylogenies, there was limited support for partitioning these other phylogenies into distinct subtrees. Using the results in Figures 2A and 3A as a guide, I selected cutoffs at which the number of subtrees was relatively stable for each of these other segments (Supplementary Figure S4). These cutoffs tended to be around 0.05, yielding four to six subtrees (Supplementary Table S4), with the exception of the NS1/NS2 phylogeny where five subtrees were obtained at cutoffs ranging from 0.12 to 0.175. This difference in scale is consistent with previous findings that diversifying selection at the amino acid level for NS1 is comparable to the surface-exposed HA and NA proteins [4]. In every case, nearly all sequences were assigned to one of the five subtrees. Two of the remaining small subtrees corresponded to the recently described H17N10 [29] and H18N11 [30] strains, respectively. Additionally, the divergent lineage of H9N2 that was consistently separated into its own HA and NA subtrees was also separated for all other segments except PB1. For all phylogenies except NP, one of the small subtrees corresponded to isolates of equine influenza (H7N7) from 1956. Finally, the phylogeny relating concatenated M1 and M2 protein sequences contained an additional subtree comprising three reassortant sequences isolated from swine in Australia [33].

## Discussion

These results indicate that it is feasible to define subtypes on the basis of the variation among HA and NA protein sequences. Subtypes derived from phylogenies are largely consistent with the current definitions of 18 HA and 11 NA subtypes, which are mostly derived from serological data. However, there are some important discrepancies. The analysis presented here supports 17 genetically-defined HA subtypes, not 18. Two subtypes are removed by merging H15 with H7, and combining H13 and H16. This is consistent with previous work reporting average amino acid identities of 79.7% and 81.4% between these respective groups, where the typical mean identity was about 49% [29]. In addition, it supports the creation of a new subtype from a divergent lineage of H9N2-like isolates from bats. This lineage not only also creates an additional twelfth subtype for NA, but also forms distinct subtrees for nearly every other segment. The two sequences comprising this lineage (Genbank accessions MH376902 and OQ216561) were isolated from bats in Egypt in 2017 [15] and South Africa in 2018 [23], respectively. Kandeil et al. [15] reported the HA sequence was the most similar (73% nucleotide identity) to H9 sequences that circulate predominantly in avian host species. Even though this divergence was comparable to that between other HA subtypes, neither study made a case for defining a new subtype.

This is not the first attempt to cluster the genomic segments of IAV into subtypes. For instance, Lu et al. [20] performed a similar clustering analysis of roughly all segments for 2,300 complete IAV genomes. For each segment, they used neighbor-joining to reconstruct trees from the HKY85 distance matrix for the multiple sequence alignment, and then extracted clusters based on a boot-strap support threshold of *>*90% and a nucleotide p-distance threshold of about 10%. Their analysis partitioned the HA sequence phylogeny into 78 subtrees, for example. Unfortunately, the web interface for classifying IAV sequences under this system (https://www.flugenome.org) was discontinued in 2016, and the expired domain name is presently being used by a laboratory reagents supplier under the guise of the original website (last accessed June 30, 2023). In addition, the classifier developed by Lu et al. [20] was not made publicly available, and it was not described with sufficient detail to reproduce.

Similarly, Zhuang et al. [35] used neighbor joining to reconstruct phylogenies relating between 4,194 to 33,066 sequences for each of the eight IAV segments. They partitioned each tree into clusters based on the presence of a “relatively separated branch”, as well as the distribution of host species, sampling location and collection dates among tips. For example, they described three major clusters in the PB2 phylogeny. Unfortunately, their clustering method was not described with sufficient quantitative detail to reproduce, and the study data (*e.g.*, annotated trees) are not available online. Finally, there are multiple studies that have clustered sequences for all IAV genome segments for the purpose of detecting reassortment events. For example, Gong et al. [9] partitioned the maximum likelihood trees for 40,296 full IAV genomes into clusters based on the mean tip-to-tip distances associated with internal nodes, *i.e.*, the *y* statistic in this study. Using the overall mean of *y* as a cutoff for each segment, they obtained 493 to 663 clusters per phylogeny. Putative reassortment events were identified from genomes with novel combinations of clusters among segments. Although this fine-grained clustering may confer greater sensitivity for detecting reassortment, it does not provide insight into the genetic characteristics of established IAV subtypes.

The process of updating the nomenclature of IAV subtypes has been complicated by the transition from serological testing to genetic sequencing to characterize novel variants (Table 1). Given the serological origins of most IAV subtype definitions, it is remarkable that these groupings can be readily recovered from genetic variation alone, although not all methods are equally successful. There are significant advantages of developing a genetic taxonomy for IAV subtypes. The number of published IAV sequences is growing exponentially (Supplementary Figure S1). IAV infections are increasingly subtyped by genetic sequencing methods that are more scaleable than serological testing. At the current pace, we can expect nearly half a million new IAV sequences to have been deposited in GISAID by the end of 2023. We can also expect an increasing number of these sequences to be derived from environmental and metagenomic samples [10], where intact virus particles would not be available for serological characterization. Expanded sampling of non-human, non-domesticated host species with these technologies will increase our chances of encountering more novel and highly divergent lineages [7, 29, 30]. Finally, a major advantage of a genetic taxonomy is that sequence-based criteria for defining subtypes can be transparent and reproducible if the clustering methods are described sufficiently well, or better yet, made freely available online.

## Data availability

The multiple sequence alignments and maximum likelihood phylogenies for all eight IAV segments in this study are available under a Creative Commons license at https://doi.org/10.5281/ zenodo.8119571. The Python and R scripts used to produce and analyze these data are released for unrestricted use and modification under the MIT license at https://github.com/PoonLab/fluclades.

## Acknowledgements

This study was motivated and informed by discussions with Dr. Richard Neher, who read an earlier version of this manuscript and suggested using mutual information to measure the correlation between phylogenetic subtrees and subtype labels. Dr. Sarah Otto provided additional feedback on the manuscript, including some improvements to figures.

## Supplementary Figures

**Figure S1:**
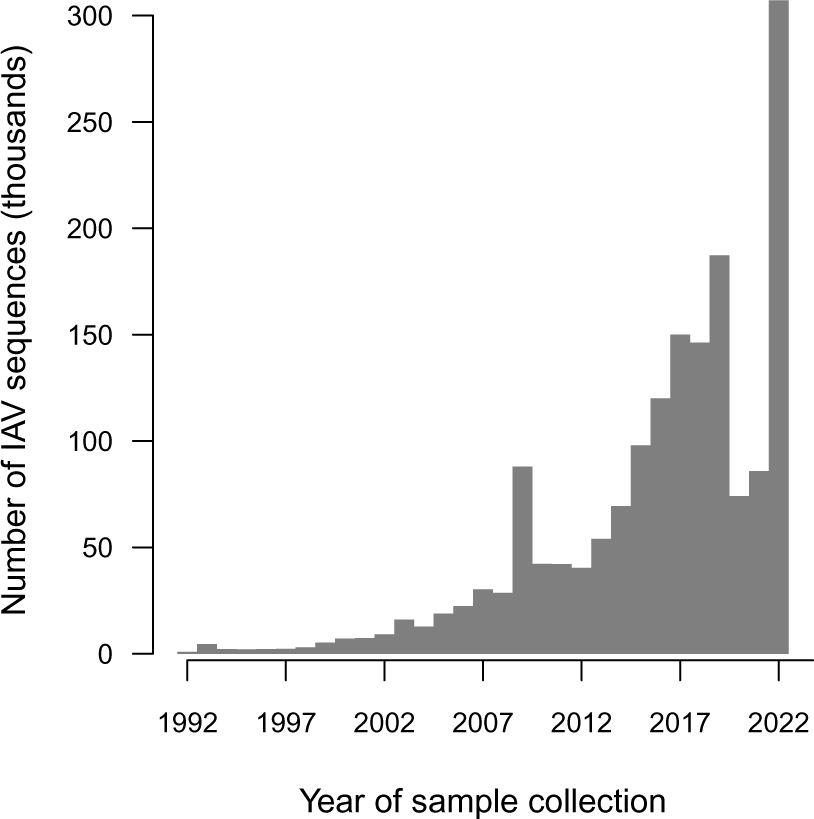
Summary of the number of influenza A virus sequences in the GISAID database by year of sample collection. Note that some sequences were collected prior to the establishment of this database in 2006, and submitted retrospectively at a later date. The number of sequences per isolate varies not only because of varying numbers of segments sequenced, but also because some isolates were sequenced more than once. This trend features a conspicuous surge in the number of sequences associated with the 2009 H1N1 pandemic, as well a drop in numbers in 2019-2020 in association with the SARS-CoV-2 pandemic. Overall, the number of sequences is exponentially growing over time (adjusted *R*^2^= 0.926).

**Figure S2:**
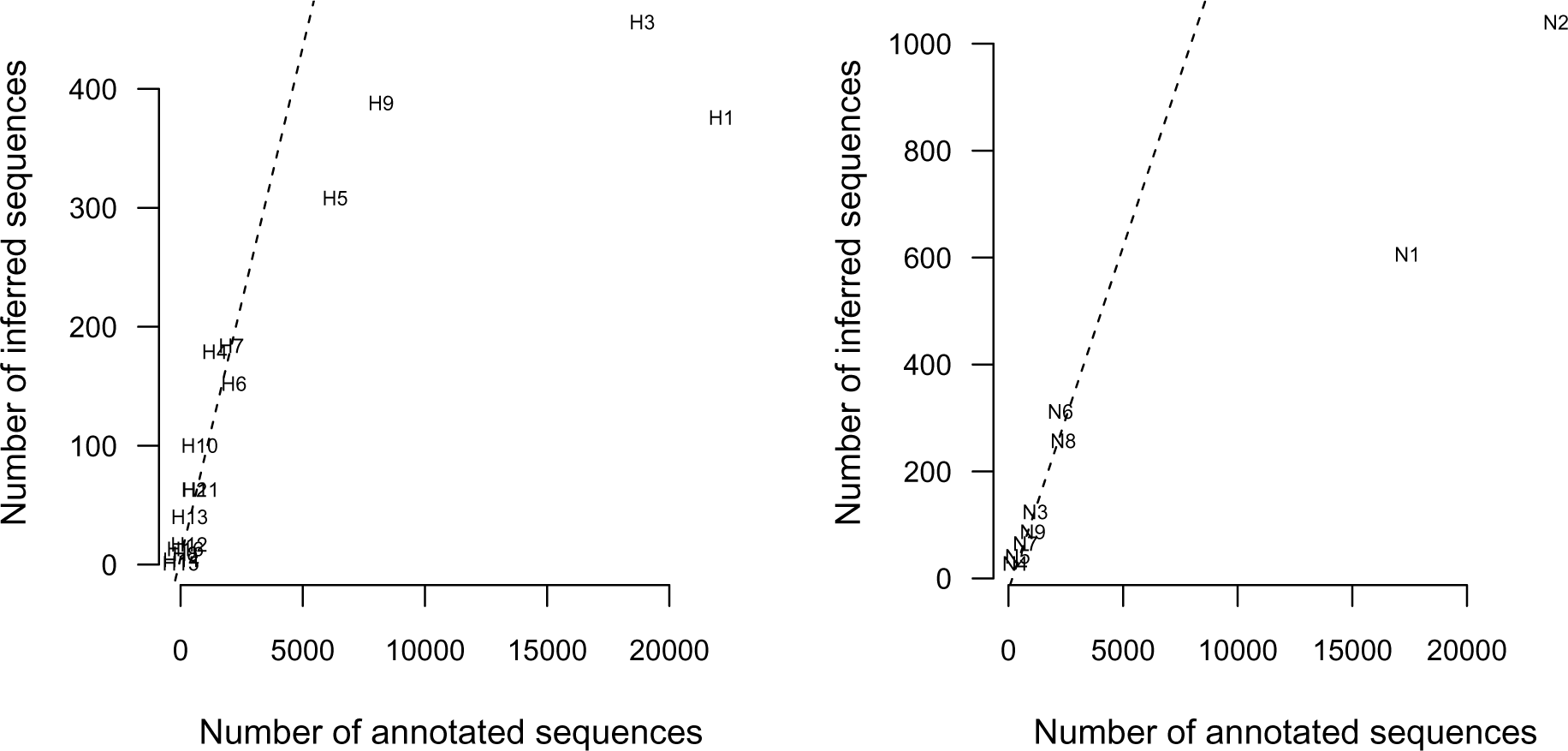
Inferred subtypes for HA (left) and NA(right) sequences with missing subtype annotations (*e.g.*, ‘unknown’, ‘HX’ or ‘mixed’) compared to their respective known distributions. Subtypes were inferred by retrieving each sequence’s nearest neighbour in the phylogeny. Dashed lines represent linear regressions on counts excluding the most common subtypes (H1, H3, H5 and H9 for hemagglutinin; N1 and N2 for neuraminidase).

**Figure S3:**
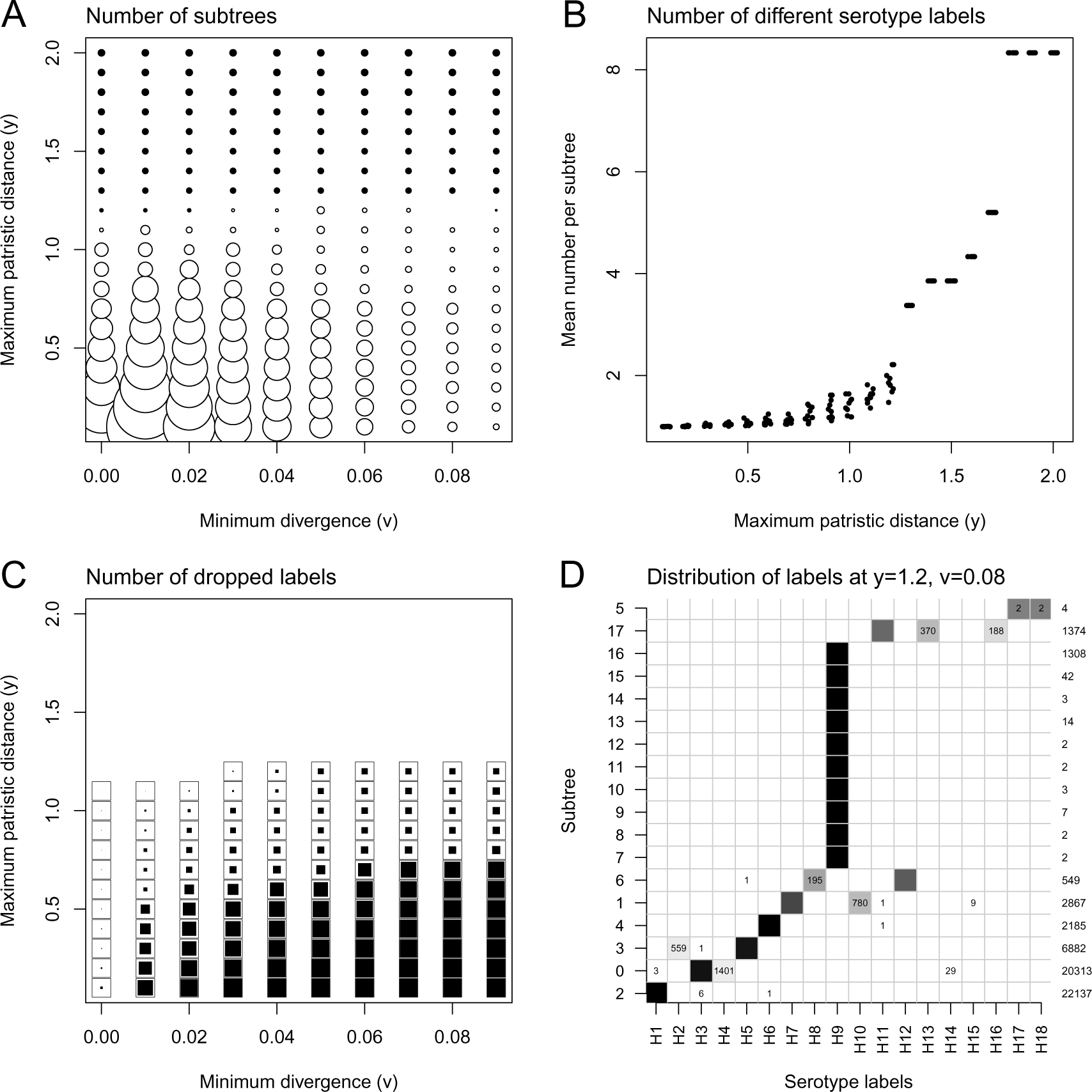
Evaluation of concordance between HA subtype annotations (labels) and subtrees under varying thresholds of nodewise clustering, for comparison to edgewise clustering results summarized in Figure 2. (A) The size of each point is scaled to the discordance between the number of subtrees and the target number (*n* = 18), *i.e.*, smaller is better. Open circles indicate too many subtrees, and filled circles indicate too few. (B) The number of different labels increases as a function of the mean patristic distance cutoff (*y*). The ideal number is one label per subtree. Varying the cutoff on minimum divergence (*v*) had a limited effect on this outcome. (C) The proportion of labels that are not associated with any subtree at given cutoffs are represented by filled squares. For reference, an outline is drawn for each square to represent 100% loss. (D) The distribution of labels among subtrees defined by cutoffs at which the ideal number of subtrees is obtained (*y* = 1.2 and *v* = 0.08). Regions are shaded in proportion to the fraction of each label in the subtree, and the number of labels is displayed if the fraction is *≤* 0.5. The total number of labels per subtree is displayed along the right margin.

**Figure S4:**
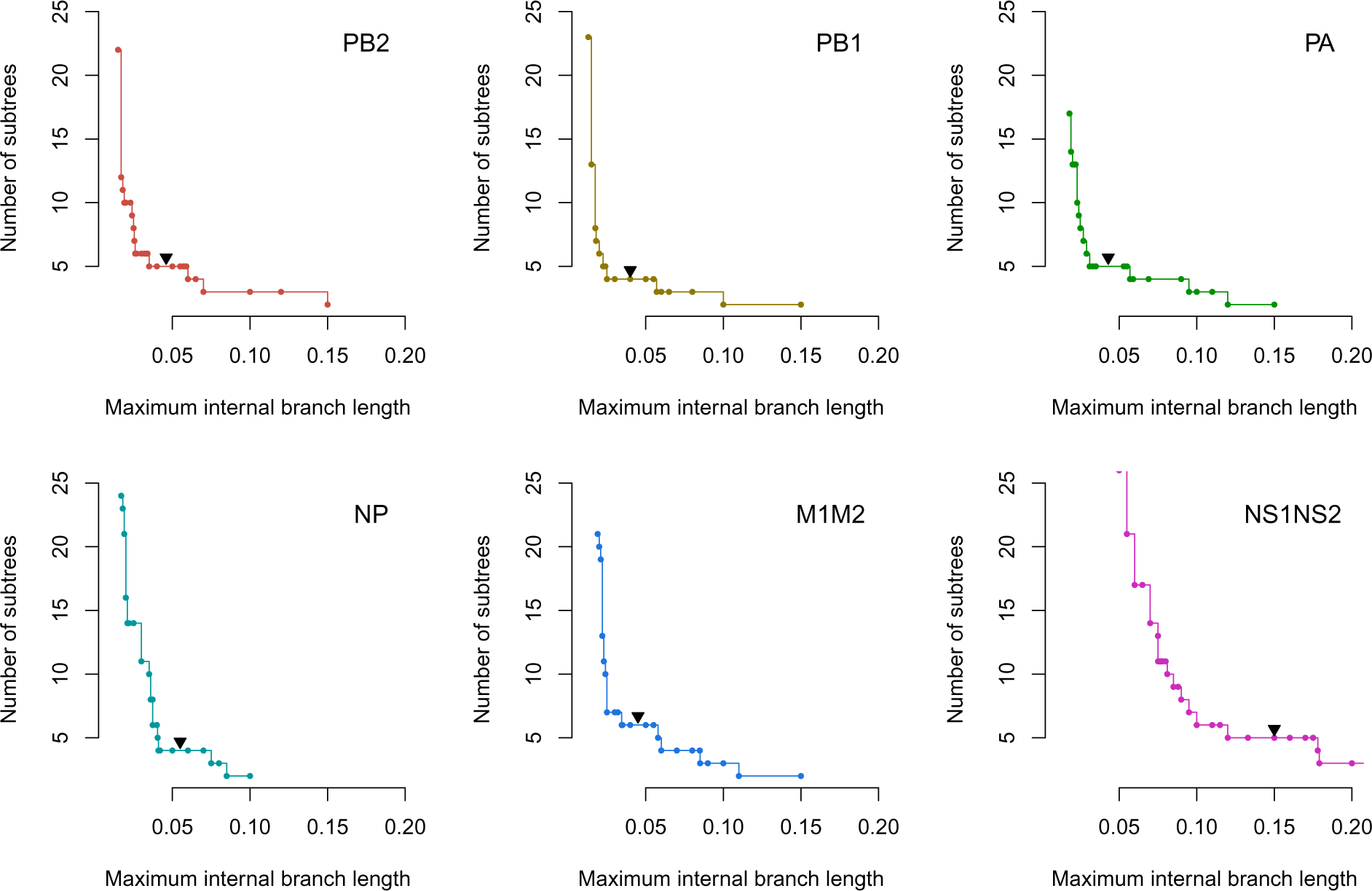
Number of subtrees as a function of internal branch length cutoffs applied to the phylogenies of other proteins. The plots are ordered with respect to genome segment numbering, *e.g.*, protein PB2 is encoded by segment 1. Solid triangles mark the cutoffs where the composition of the resulting subtrees are summarized in Supplementary Table S4. Since the edgewise clustering method yields substantially fewer subtrees for these phylogenies than HA and NA, we cannot use the same approach of taking the longest segment for the most robust number of subtrees. Instead, I manually selected the first long segment past the subjective ‘elbow’ for each trend. As shown in Supplementary Table S4, changing this cutoff has little impact on the composition of subtrees.

## Supplementary Tables

**Table S1:** Summary of amino acid sequences obtained for all eight IAV segments from the NCBI Genbank nucleotide database (date accessed, June 1, 2023 except for HA sequences, which were retrieved April 28, 2023). Amino acid sequences representing PB1-F2 and PA-X (italics) were excluded from subsequent analyses. The M1/M2 and NS1/NS2 sequences were concatenated with amino acid residues within overlapping regions excluded. Min. length = minimum sequence length in nucleotides. Total = number of sequence records returned by query. Unique = number of unique amino acid sequences after filtering.

| Segment | Gene products | Min. length | Total | Unique |
| --- | --- | --- | --- | --- |
| 1 | PB2 | 2000 | 93,415 | 32,718 |
| 2 | PB1, *PB1-F2* | 2000 | 160,506 | 29,790 |
| 3 | PA, *PA-X* | 2000 | 169,484 | 32,349 |
| 4 | HA | 1600 | 137,172 | 66,874 |
| 5 | NP | 1400 | 103,033 | 18,978 |
| 6 | NA | 1000 | 126,233 | 52,146 |
| 7 | M1, M2 | 900 | 213,717 | 14,389 |
| 8 | NS1, NS2 | 750 | 191,676 | 25,172 |

**Table S2:** Reclassification of IAV HA sequences with serotype (subtype) annotations that are not consistent with their location in the phylogeny. Discordant sequences were detected as outliers in box-and-whisker plots of their vertical positions in the rectangular layout of the tree, grouped by subtype annotations. The predicted subtypes were estimated from the nearest labeled tips in the tree. These predictions were manually verified using nucleotide BLAST.

| Accession | Strain | Original | Corrected |
| --- | --- | --- | --- |
| ON427060 | A/swine/Kansas/A02711700/2022 | H1 | H3 |
| MF098827 | A/swine/Chile/VN1401-587/2014 | H1 | H3 |
| MT377993 | A/swine/Thailand/CU22337/2018 | H1 | H3 |
| MH596892 | A/ruddy turnstone/Delaware/650622/2002 | H11 | H7 |
| MN569599 | A/Newcastle/177/2017 | H3 | H1 |
| MN574095 | A/Victoria/79/2017 | H3 | H1 |
| KM110063 | A/swine/Hanoi/568/2013 | H3 | H1 |
| KM110062 | A/swine/Hanoi/424/2013 | H3 | H1 |
| KM110061 | A/swine/Hanoi/411/2013 | H3 | H1 |
| CY251407 | A/Victoria/1003A/2012 | H3 | H1 |
| OP888980 | A/yellow-billed | H6 | H1 |
| LC500374 | A/duck/Vietnam/HU8-1918/2017 | H3 | H5 |
| MH598081 | A/ruddy turnstone/New Jersey/650677/2002 | H11 | H6 |
| MH598105 | A/ruddy turnstone/New Jersey/828212/2001 | H5 | H12 |
| MH130194 | A/mallard/Korea/M198/2013 | H4 | H9 |

**Table S3:** Reclassification of IAV NA sequences with serotype (subtype) annotations using the same methods described in Table S2.

| Accession | Strain | Original | Corrected |
| --- | --- | --- | --- |
| MK960206 | A/Texas/9437/2019 | N1 | N2 |
| OP940105 | A/swine/Iowa/G92995/2022 | N1 | N2 |
| OP645900 | A/swine/Iowa/C61728/2022 | N1 | N2 |
| OP940090 | A/swine/Illinois/E80174/2022 | N1 | N2 |
| MW839927 | A/swine/Iowa/CD81015/2015 | N1 | N2 |
| MW839656 | A/swine/North Carolina/LJ8316/2016 | N1 | N2 |
| MT377995 | A/swine/Thailand/CU22337/2018 | N1 | N2 |
| MF099208 | A/swine/Chile/VN1401-587/2014 | N1 | N2 |
| MT378776 | A/swine/France/22-180459/2018 | N1 | N2 |
| MW362711 | A/swine/Belgium/Gent-36/2016 | N1 | N2 |
| MT818840 | A/Mallard/North Carolina/AH0114261I.6.A/2017 | N8 | N2 |
| MW513533 | A/green-winged teal/Alaska/UGAI16-4602/2016 | N8 | N2 |
| MF613906 | A/ruddy turnstone/Delaware Bay/303/2016 | N4 | N2 |
| MH597119 | A/ruddy turnstone/Delaware/1016391/2003 | N5 | N2 |
| MH597724 | A/ruddy turnstone/New Jersey/1321395/2005 | N8 | N6 |
| MH597785 | A/ruddy turnstone/New Jersey/1321408B/2005 | N8 | N6 |
| MH598073 | A/ruddy turnstone/New Jersey/650675/2002 | N9 | N5 |
| MH597447 | A/ruddy turnstone/Delaware/650647/2002 | N9 | N5 |
| L06581 | A/equine/New Market/1979 | N3 | N8 |
| MH597692 | A/ruddy turnstone/New Jersey/1148671/2004 | N7 | N8 |
| MH596859 | A/ruddy turnstone/Delaware/1016396/2003 | N9 | N8 |
| LC431470 | A/pintail/Alberta/121/1979 | N8 | N4 |
| CY251409 | A/Victoria/1003A/2012 | N2 | N1 |
| MH598004 | A/ruddy turnstone/New Jersey/650584/2002 | N9 | N1 |
| MH596894 | A/ruddy turnstone/Delaware/650622/2002 | N4 | N1 |
| MH579375 | A/wild waterfowl/Korea/M54-F-1/2010 | N2 | N1 |
| MK636253 | A/New York/7745/2018 | N2 | N1 |
| ON848942 | A/Homosapiens/India/149/2017 | N2 | N1 |
| MN568417 | A/Brisbane/144/2017 | N2 | N1 |
| MN574097 | A/Victoria/79/2017 | N2 | N1 |

**Table S4:**
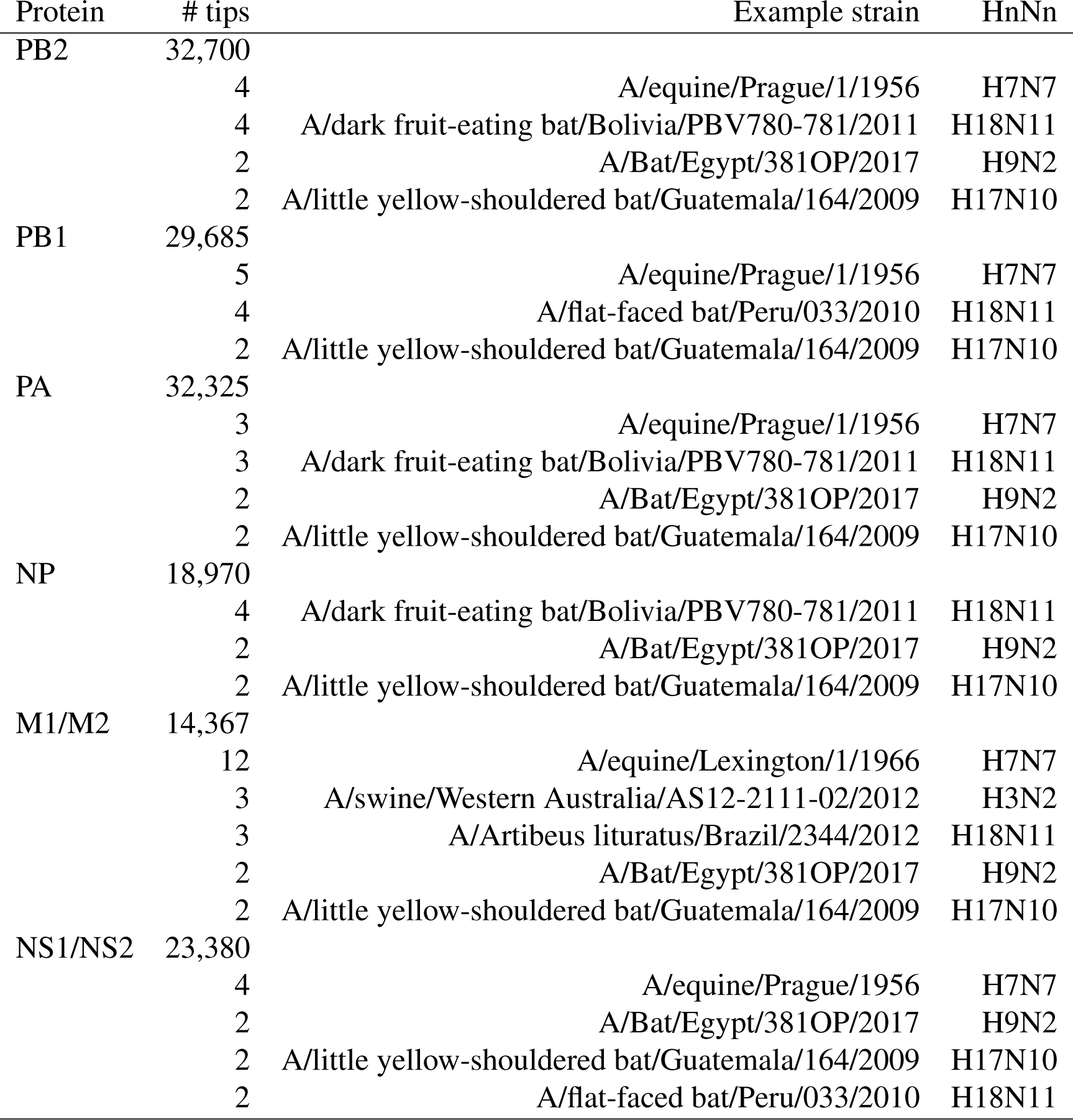
Composition of subtrees produced at selected branch length cutoffs for phylogenies relating other protein sequences. Cutoffs were chosen based on the trends displayed in Supplementary Figure S4. Example strain and HA/NA subtype labels (HnNn) are omitted for the largest subtrees.

## Notes

https://doi.org/10.5281/zenodo.8119571

https://github.com/PoonLab/fluclades/

### Competing Interest Statement

The authors have declared no competing interest.

### Summary of Updates

Added normalized mutual information results to text; revised Figures 2 and 3 for clarity; fixed some typographical errors; some modifications to supplementary figures S1 and S3 for clarity; provided more explanation of selecting cutoffs for Supplementary Figure S4.

